# Transformation of the symbiotic alga Oophila amblystomatis : a new toolbox for animal-algae symbiosis studies

**DOI:** 10.1101/2022.04.06.487339

**Authors:** Baptiste Genot, John A Burns

**Affiliations:** Bigelow Laboratory for Ocean Sciences, East Boothbay, ME

**Keywords:** symbiosis, algae, transformation

## Abstract

The ability to conduct reverse genetic studies in symbiotic systems is enabled by transgene expression and transformation of at least one partner. The symbiotic relationship between the yellow spotted salamander, *Ambystoma maculatum*, and the green alga, *Oophila amblystomatis*, is a unique model of vertebrate-algae symbiosis. Despite over 130 years of scientific study, there are still many open questions in this symbiosis. Transgene expression in one partner will accelerate research into the symbiotic relationship. In this paper we describe a tool and method for expression of foreign DNA in, and presumed transformation of, the alga *O. amblystomatis*. We successfully introduced heritable antibiotic resistance to algal cultures, and observed expression of a green fluorescent reporter protein in all transfected and presumably transformed algal populations. The outcomes of this work enable genetic manipulation of the symbiotic alga *Oophila amblystomatis*, allowing direct testing of hypotheses derived from gene expression or genomic studies that will usher in a deeper understanding of the *A. maculatum*-*O. amblystomatis* symbiotic system.

**Summary statement:** Genetic tools stimulate new possibilities for research in living systems. This work describes a new tool for transformation of a symbiotic alga that enters vertebrate tissues and cells.

## Introduction

Symbiotic relationships, intimate interactions between two or more organisms, are common in living systems and have long intrigued the scientific community. Symbioses between heterotrophic hosts and photosynthetic algae, known as photosymbioses, are mainly known from invertebrate animals and protists as hosts paired with-algal symbionts. Iconic examples include cnidarians (reef corals, jellyfish, sea anemones, hydras) and their *zooxanthellae* (reviewed in Davy et al. 2012), acantharian and radiolarian protists and their haptophyte and dinoflagellate endosymbionts (Decelle et al. 2015), *Paramecium bursaria* and the green alga chlorella (Karakashian 1975), or lichens as fungus-alga associations (Ahmadjian 1993). Photosymbioses with vertebrate hosts are not as well known (recently reviewed in Yang et al. 2022) and only one example of a vertebrate-alga endosymbiosis has been observed between the salamander *Ambystoma maculatum* and the green alga *Oophila amblystomatis*. Green algae in the *Oophila* clade also enter facultative symbiotic relationships with the eggs and embryos of a variety of amphibians (Kim et al. 2014).

Important advances in biological research have come from analyses of organismal systems using forward and reverse genetics. With the advent of sequencing based research, targeted reverse genetics has become a powerful tool to test hypotheses about gene function in many systems. Transfection and transformation protocols that allow researchers to tag and modify specific genes in model species like *Eschericia coli, Drosophila melanogaster, Caenorhabditis elegans, Arabidopsis thaliana, Danio rerio*, and *Chlamydomonas reinhardtii*, to name a few, have transformed our understanding of how cells and biological systems work. In photosymbiotic systems, reverse genetics tools are in their infancy. Many photosymbioses are currently genetically intractable in both host and symbiont. Key advances have been made in corals (Cleves et al. 2018) but while there has been success in transforming a dinoflagellate (Nimmo et al. 2019), reports of genetic manipulation in the symbiotic *Symbiodinium* clade remain controversial (Nimmo et al. 2019) and not yet widely adopted.

With available physiological and genetic data for the *A. maculatum-O. amblystomatis* symbiosis, genetic manipulation of the partners is one avenue to accelerate research into their unique partnership. The alga *O. amblystomatis* is a chlamydomonad alga, superficially similar to the model alga *C. reinhardtii*. DNA constructs for transgene expression and transformation of *C. reinhardtii* are readily available, however divergence time between *C. reinhardtii* and *O. amblystomatis* is long, for example the two green algae likely have a divergence time longer than that between humans and fish (Munakata et al. 2016). In this paper we show how we have adapted constructs and methods that are successful in transforming wild-type *C. reinhardtii* cells to the symbiotic alga *O. amblystomatis*. The use of native promoter regions from *O. amblystomatis* was essential for success. The ability to express transgenes in *O. amblystomatis* will open new avenues of research in amphibian-alga symbioses.

## Results

### Gene expression informed promoter selection

A combination of gene expression and whole genome data was used to design native promoter DNA constructs to drive transgene expression in *O. amblystomatis. Oophila* algae live in multiple habitats, including pondwater (Bishop et al. 2021), inside amphibian eggs (Kim et al. 2014), and within A. *maculatum* tissues and cells (Kerney 2011). In order to find promoters that would drive expression across known physiological conditions, gene expression data from prior studies (Burns et al. 2017; Kerney et al. 2019) was analyzed to find transcripts that were expressed at a high level and with a low variance across multiple conditions relevant to the symbiosis including cultured algae, algae from salamander egg capsules, and intracellular algae (Figure 1). A 2kbp region upstream of the start codon of candidate genes was recovered from the *O. amblystomatis* genome assembly. Alignments of the regions upstream of the presumed coding sequence in *O. amblystomatis* and related organisms were used to refine potential promoter regions (Figure S1). This led to the selection of three promoters that were cloned into plasmid constructs.

**Figure 1.**
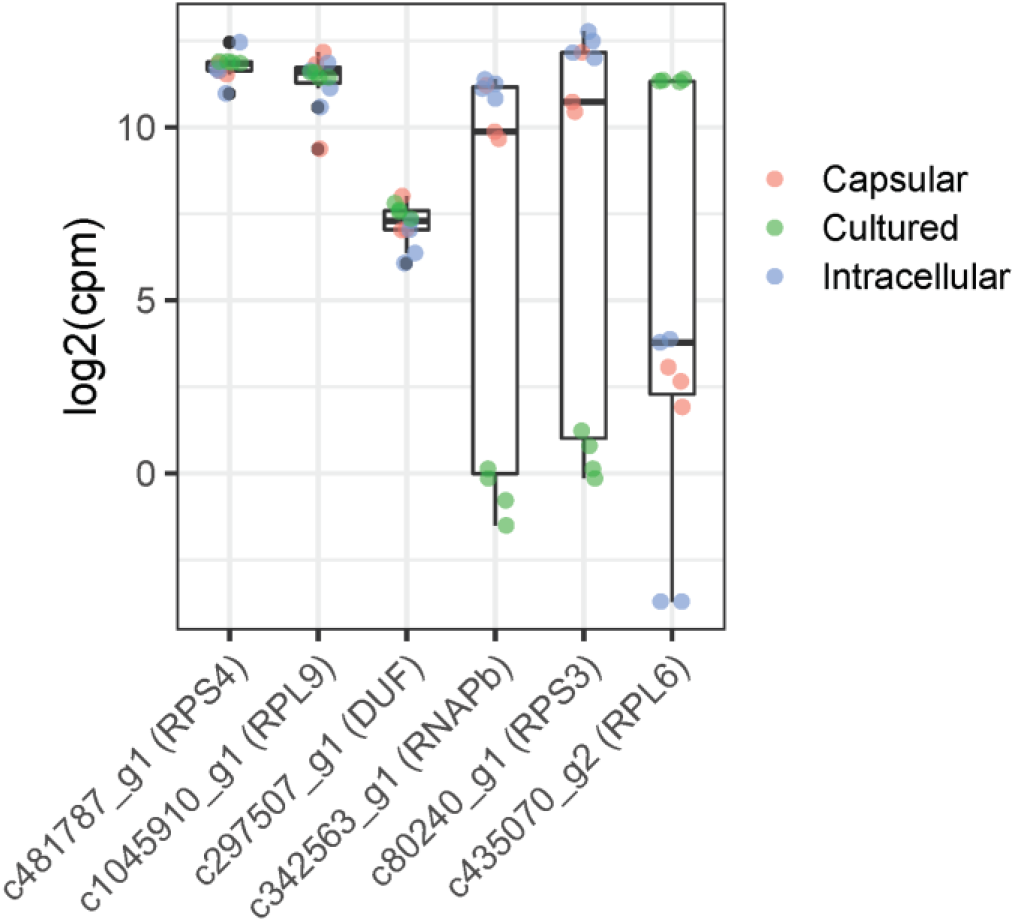
Gene expression levels under symbiotic and lab culture conditions. Upstream promoter regions were obtained for three genes with high expression levels and low variance across conditions (RPS4, RPL9, DUF). For comparison, three genes with highly variable and condition dependent expression levels are shown (RNAPb, RPS3, RPL6). Data were collected from Burns et al. 2017 and Kerney et al. 2019 and include transcripts level of capsular algae (pink color), intracellular algae (blue color) and cultured algae (green color).

For reporter and antibiotic resistance gene expression, we used the architecture of plasmid pMO508 (Onishi and Pringle 2016), optimized for transgene expression in *C. reinhardtii*. Plasmid pMO508 contains a *C. reinhardtii* promoter that drives expression of a bicistronic cassette encoding a reporter gene, the green fluorescent protein CrVENUS, and an antibiotic resistance gene, *APHVII*, on a single transcript. The symbiotic alga *O. amblystomatis* has similar codon usage to *C. reinhardtii* (Figure S2), suggesting the *C. reinhardtii* optimized coding sequences could be transcribed and translated efficiently in *O. amblystomatis*. Wild type *O. amblystomatis* cultures were found to be naturally sensitive to the antibiotic paromomycin (Figure S3), suggesting that expression of the gene *APHVIII* would allow selection of transformants while expression of CrVENUS would allow visualization of transgene expression.

To test native promoter efficacy for transgene expression, we modified pMO508 by replacing the *C. reinhardtii* pTUB2 promoter and adjacent RbSC2 intron with native *O. amblystomatis* promoter regions (Figure 2, Table 1). We also created a construct with an *O. amblystomatis* promoter and the *Chlamydomonas* RbSC2 intron between the promoter and the ATG of the *CrVENUS* gene, to see whether the intron could enhance transgene expression as has been observed in other systems (Baier et al. 2018; Suttangkakul et al. 2019).

**Figure 2.**
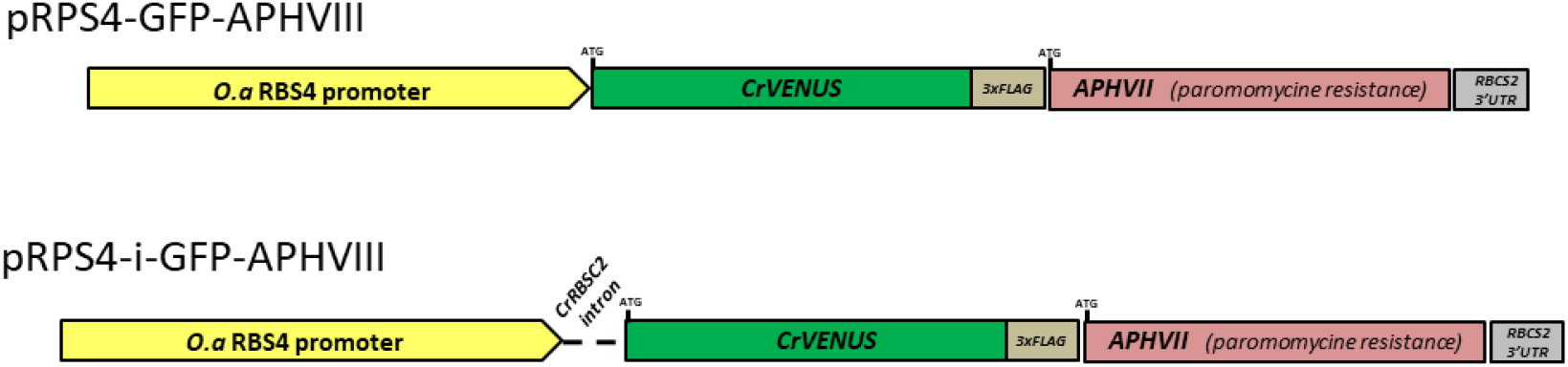
Schematic representation of the plasmid constructs created for *O. amblystomatis* transfection.

**Table 1:** Summary of plasmids, and transfection outcomes, used for transfection and transformation of *O. amblystomatis*.

| Plasmid | Reference | Optimized for | Source/backbone | Paromomycin resistance |
| --- | --- | --- | --- | --- |
| pM0508<br>pTUB-i-GFP-APHVII | Onishi and Pringle 2016 | *Chlamydomonas reinhardtii* | chlamycollection.org | no |
| pM0507<br>pPSAD-i-GFP-APHVII | Onishi and Pringle 2016 | *Chlamydomonas reinhardtii* | chlamycollection.org | no |
| pCaMV35S-i-GFP-APHVII | Not published | Land plants | pMO508 | no |
| p_c1045910-i-GFP-APHVIII | Not published | *Oophila amblystomatis* | pMO508 | no |
| p_c297507-i-GFP-APHVIII | Not published | *Oophila amblystomatis* | pMO508 | no |
| pRPS4-i-GFP-APHVIII | This paper | *Oophila amblystomatis* | pMO508 | yes |
| pRPS4-GFP-APHVIII | This paper | *Oophila amblystomatis* | pMO508 | yes |

We also tested the un-modified base plasmids, pMO508 and pMO507 (Onishi and Pringle 2016) to see whether the *C. reinhardtii* promoters would drive transgene expression. In addition, a pMO508-derived plasmid with the plant Cauliflower Mosaic Virus (CaMV) 35S promoter cut from a pCAMBIA1302 (pCaMV35S-i-GFP-APHVII) was created since the 35S promoter has successfully driven expression of reporter genes in other unicellular organisms (Faktorová et al. 2020) (Table 1).

### The *O. amblystomatis* Ribosomal Protein S4 promoter drives transgene expression and transformation of the alga

Electroporation with linearized plasmid that used the upstream region of the Ribosomal Protein S4 (RPS4) gene to drive transgene expression consistently produced paromomycin resistant cultures (Table 1, Figure S4 for sequence data). Plasmids with other promoters failed to induce transformation of *O. amblystomatis* (Table 1). Complementary methods were used to detect transgene expression in paromomycin resistant cultures. RT-qPCR detected transgene mRNA, western blots detected protein products produced from the transgene mRNA, and microscopy and flow cytometry were used to visualize and quantify reporter gene (*CrVENUS*) activity.

Transgene mRNA was amplified in all antibiotic-resistant cultures while it was not detected in wild type samples. Transgene expression is highly variable between transformed lines derived from independent transfection trials. No significant differences in mRNA quantity were observed between lines transformed with plasmids carrying the CrRBSC2 intron and lines transformed with plasmids that did not have the intron, suggesting that the presence of the intron does not impact transgenic mRNA expression.

A band at 30kDa in western blot experiments (Figure 3) confirmed that paromomycin resistant cultures produced the FLAG-tagged reporter protein (CrVENUS-FLAG) and that it appeared at the expected size of 30 kDa (Figure 3), indicating that the bicistronic mRNA codes for production of two separate protein products as intended. Additionally, the relative abundance of CrVENUS-FLAG protein correlates with transcript abundance for each independently derived transformed line. The transformed line pRPS4-i-GFP-APHVIII(A) has the highest mRNA expression level of the transgene, and has the most intense band for the CrVENUS-FLAG protein (Figures 4 and 3, respectively). On average, however, there does not appear to be a difference between transformed lines that carry the CrRBSC2 intron and those that do not.

**Figure 3.**
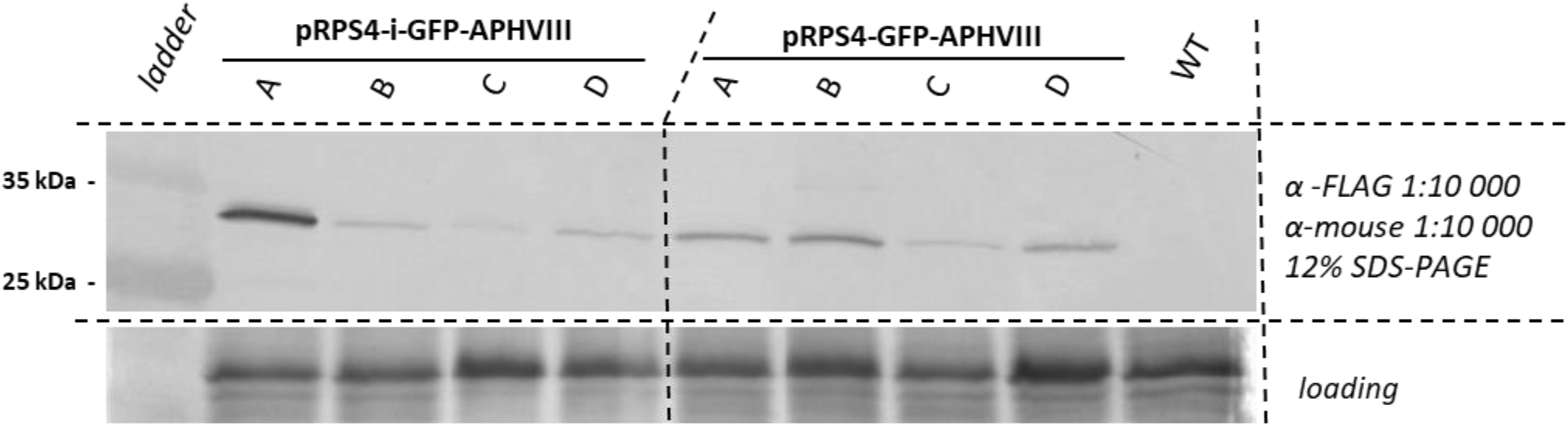
Paromomycin resistant algal cultures make CrVENUS-FLAG protein. Western blot comparing CrVENUS-FLAG protein levels (α-FLAG primary antibody) in cultures derived from independent transfections. 10 µg total proteins were loaded in each well and resolved via SDS-PAGE before blotting. The loading control is a coomassie stain of total protein remaining in the gel after blotting.

**Figure 4.**
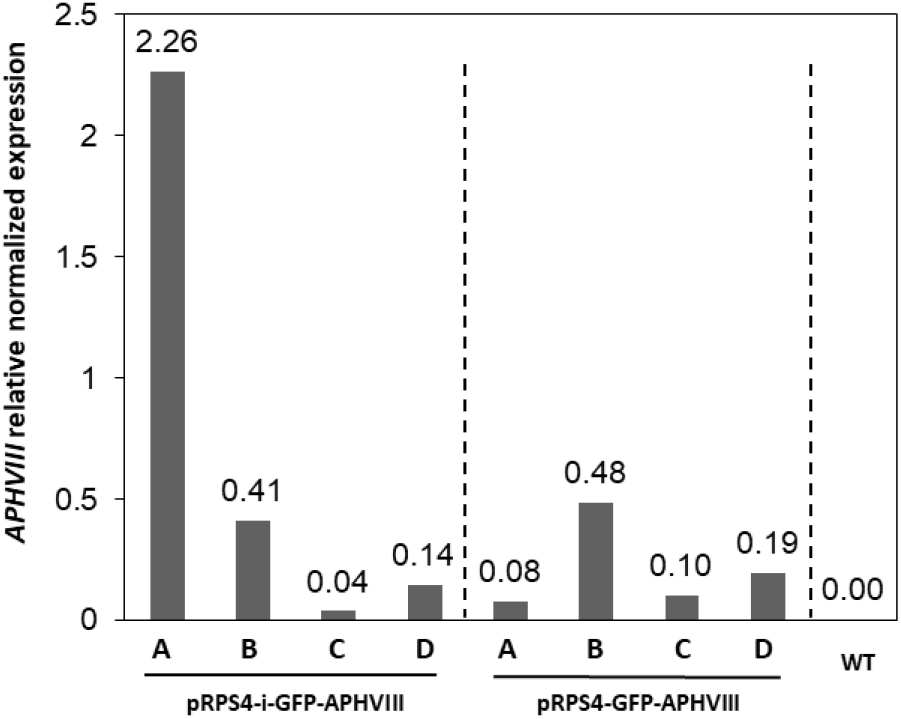
Paromomycin resistant algal cultures express plasmid specific mRNA. RT-qPCR reactions used primers targeting the *APHVII* coding region (near the 3′ end) of the bicistronic, plasmid derived transcript. No statistically significant differences were found between plasmids with intron and plasmid without intron ANOVA). Transcripts derived from untransformed WT cells had no amplification using *APHVII* specific primers.

Reporter gene activity in paromomycin resistant cultures was confirmed using fluorescence microscopy and flow cytometry. The two transformed lines that exhibited the highest mRNA and protein levels, pRPS4-i-GFP-APHVIII(A) and pRPS4-GFP-APHVIII(B), also displayed visible GFP fluorescence by fluorescence microscopy (Ex 470 nm/Em 590 nm) (Figure 5). However, while GFP fluorescence could be observed in both transformed lines, fluorescence was highly variable on a cell-by-cell basis and a large proportion of cells had a weak GFP signal. Flow cytometric quantification of GFP fluorescence in all transformed lines showed that all paromomycin resistant cultures exhibited green fluorescence in excess of the background signal from untransformed, wild type *O. amblystomatis* cells (Figure 5). In line with RT-qPCR and western blot data, transformed line pRPS4-i-GFP-APHVIII(A) showed the highest average GFP fluorescence. Similarly, in line with other data, pRPS4-GFP-APHVIII(B) showed the highest average GFP fluorescence among lines transformed with the plasmid lacking the RBSC2 intron.

**Figure 5.**
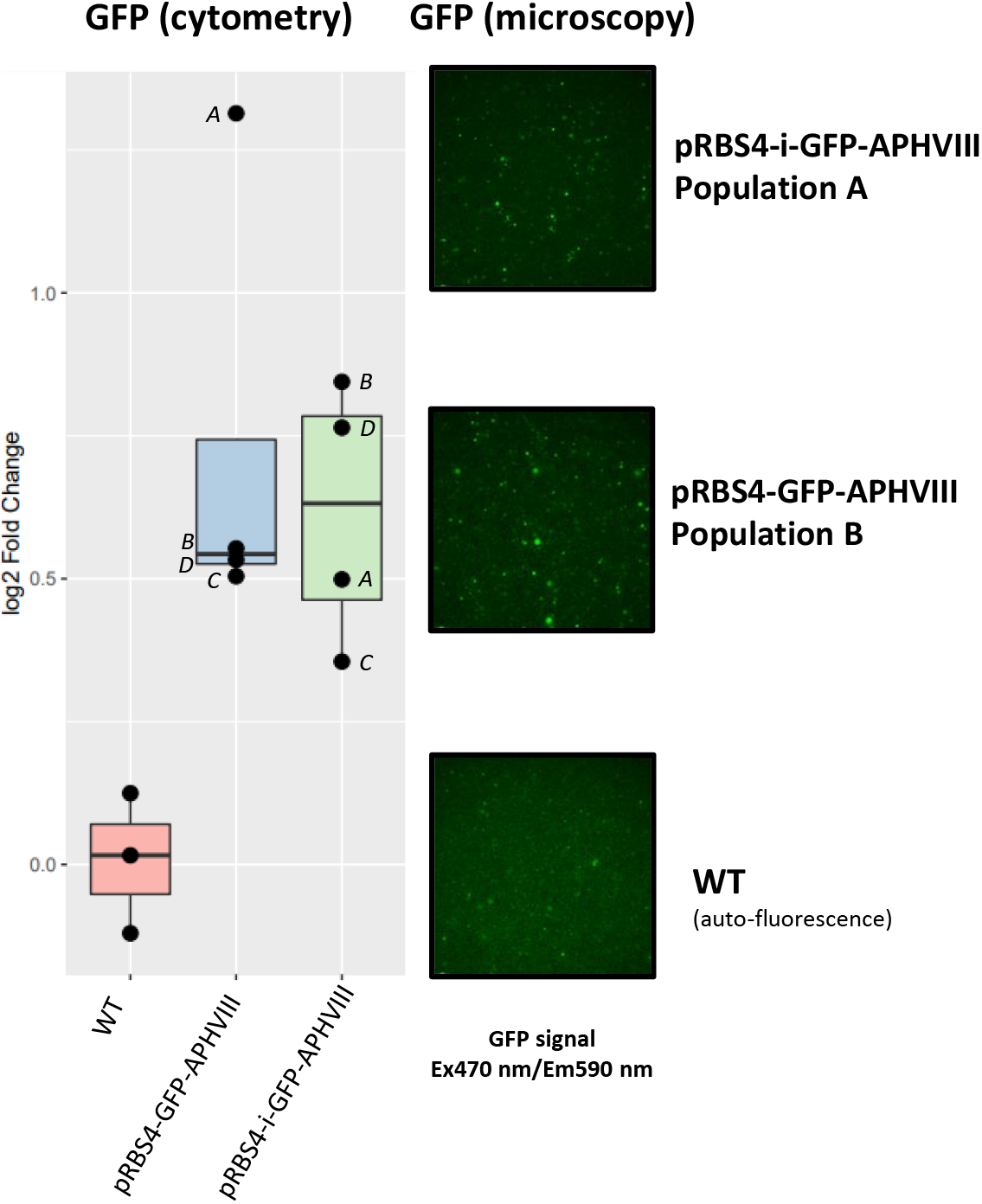
Paromomycin resistant algal cultures show specific green fluorescence of the CrVENUS protein. Left panel: GFP fluorescence of all transfected lines quantified by flow cytometry in comparison to wild type *O. amblystomatis*. Values can be interpreted as log_2_(fold-change) in fluorescence intensity with respect to the median GFP fluorescence in WT cells, calculated as log_2_(sample GFP) - median(log_2_(WT GFP)). Right panel: Photomicrographs showing GFP fluorescence from the two cultures with the highest transgenic protein and mRNA levels from intron containing and intron lacking constructs in comparison to wild type *O. amblystomatis*.

## Discussion

In this paper we describe a new and reliable method to express foreign DNA in the symbiotic alga *O. amblystomatis*. The algal strain that we transformed has no specific modifications with respect to wild-type *O. amblystomatis*. It contains a cell wall and is competent for invasion of host salamanders (Kerney et al. 2019). We show expression of transgene mRNA and protein accumulation from eight independent transfection trials in addition to showing transgenic protein function by way of antibiotic resistance and GFP fluorescence.

A limitation of the mixed culture selection method reported here is that while we were able to create paromomycin-resistant populations of *O. amblystomatis*, the resultant cultures do not originate from a single cell and are therefore not isogenic. *O. amblystomatis* is resistant to culture on agar and does not tend to form colonies from single cells (Kim et al. 2014), so other methods need to be used to reduce or eliminate heterogeneity of transformants, such as dilution to extinction of mixed transformed cultures or selection by cell sorting followed by culturing. Future isolation of isogenic strains of *Oophila* would allow more detailed studies regarding transgene expression and transgene copy number. Such efforts were beyond the scope of this proof of principle study on genetic constructs and methods of transformation.

Researchers are making continual progress on transfection, transformation, and engineering of non-model protists (Faktorová et al. 2020) in an effort to build a more comprehensive view of molecular and cell biology across life’s diversity. For example, in recent work transgenic expression of YFP fusion proteins in the chlorarachniophyte *Amorphochlora amoebiformis* was applied to verify the subcellular localization of an uncharacterized protein group of carbonic anhydrases (Hirakawa et al. 2021) and two DNA polymerases (Hirakawa and Watanabe 2019).

In photosymbioses, cnidarian hosts have been manipulated by transgene expression (Jones et al. 2018), RNA interference (RNAi)(Dunn et al. 2007), and gene editing tools (Cleves et al. 2018), some of which have begun to aid molecular and cellular characterization of symbiont bearing cells (Jones et al. 2018). In algal symbionts, progress is being made in transfection of dinoflagellate and haptophyte algae (Endo et al. 2016; Nimmo et al. 2019) that may lead to new insights into important animal-alga and protist-alga symbioses like those of corals and acantharians, respectively. Transfection and transformation of green algae is established for model species like *C. reinhardtii* and *Chlorella vulgaris*. Until now, transfection and transformation of a symbiotic green alga has not been demonstrated. This new electroporation and transformation protocol for *O. amblystomatis* that uses a native promoter that may permit stable transgene expression across symbiotic states will enable a greater understanding of alga-amphibian symbioses. From this starting point, future work can make use of the suite of reverse-genetic tools including knock-ins, knock-downs, and precise editing of the *O. amblystomatis* genome using the CRISPR-cas9 system.

## Figures list

**Figure S1**: Promoter selection using alignments of genomic DNA and coding sequences of target genes from related species.

**Figure S2**: *O. amblystomatis* codon usage compared to *C. reinhardtii*.

**Figure S3**: *Oophila amblystomatis* cells are sensitive to paromomycin.

*Oophila amblystomatis* cultures in AF6 media 7 days after treatment with 20 and 40 µg/mL paromomycin.

**Figure S4**: Plasmid sequences (.ape files) created for this study. Sequenced regions are in upper case.

## Materials & Methods

### Algae strain and culture

*Oophila amblystomatis* UTEX-LB-3005 was used for this study. Cells were cultivated in a modified AF6 media at 16 °C, 12 hours daylight (170 µE/m2/s) in a Percival culture chamber. Modified AF6 media was made without ammonia but with 3.85mM NaNO3 and trace metals from Kropat et al. 2011.

### Genomic DNA extraction from Oophilla amblystomatis cells

Quick genomic DNA extraction from *Oophilla amblystomatis* cells was carried out using a modified Eukaryotic Microalgal Nucleic Acids Extraction (EMN) protocol from Kim et al. 2012. 5 mL of algae from a fully grown liquid culture were pelleted at 400g for 5 min and re-suspended in 800µL h20 and 600µL phenol (pH 8). 0.5g of 0.5 mm zirconia/silica beads and 0.4g of 0.1 mm glass beads were added and samples were shaken in a Tissue lyser at 25 Hz for 5 minutes. Following cell lysis with glass beads, debris were pelleted in a microcentrifuge at >10,000g. The aqueous layer was recovered and one volume of chloroform was added to the tube. After another 5 minutes spin at >10,000g the aqueous layer was recovered and one volume of isopropanol and 1/10 volume of 3M Sodium Acetate (pH 8) were added. DNA was pelleted by centrifugation for 45 minutes at >10,000 g and washed in 70 % Ethanol before being air-dried and re-suspended in TE buffer (10mM Tris, 0.1mM EDTA, pH 7.5).

### Cloning Oophila amblystomatis pRPS4 in a Chlamydomomas expression vector

pMO508 (pTUB2_CrVenus-3FLAG) created by Onishi and Pringle 2016 was purchased from Chlamydomonas Resource Center (www.chlamycollection.org). The original pTUB2 promoter and the RbSC2 intron were cut out of the backbone using *HpaI* and *HindIII* restriction enzymes (Thermo Fast-digest) and, after checking the digest using agarose gel electrophoresis, the product of that digestion was used directly for Gibson assembly reactions.

### pRPS4-GFP-APHVIII plasmid

A long genomic DNA fragment containing *Oophila amblystomatis* pRPS4 was amplified using a nested primer approach. Outside primers p107 and p108 were used in the initial amplification from genomic DNA and inside primers p117 and p207 were used to amplify the region of interest from the initial amplicon. The inside primers included 23 bp overlaps with the destination plasmid on each side of the amplicon. Gibson assembly reaction (GeneArt Gibson Assembly HiFi, Invitrogen) was carried out according to the manufacturer’s instructions.

### pRPS4-i-GFP-APHVIII plasmid

The RbSC2 intron was amplified using p212 and p213 and *Oophila amblystomatis* promoter pRPS4 with p117 p208. The two fragments were assembled in the *HpaI*/*HindIII* pMO508 using a Gibson assembly reaction.

For both constructs, single *E*.*coli* colonies were picked up and checked by Sanger sequencing to confirm the presence of the integrity of the promoter. Prior to electroporation of plasmids into algal cells, 15 µg of plasmid DNA was linearized using the enzyme *Kpn1* (Thermo Fast-digest). See Figure S4 for plasmid sequences.

#### Primer list

p107CTCACCACTTGATAGCGTCGTCCTTGGTGACCTTG;

p108 GTGACCCACTTGAGCCTGTTCCTCAGAATC ;

p117tgccgggagcagacaagatatcaCTCTCCAGGTTGACGTAGAAGTC;

p207tcctcgcccttgctcaccatgttGCAAACAAGCCTCTAGGCAGTTGCAAG;

p208 GCAAACAAGCCTCTAGGCAGTTGCAAG ;

p212CTTGCAACTGCCTAGAGGCTTGTTTGCgccaggtgagtcgacgagcaag;

p213 tcctcgcccttgctcaccatgttAACCTCGAATCTCCTGCAAATGGAAAC

#### Electroporation

100 mL of culture grown to maximum density (∼10^6^ cells/ mL) were pelleted (5 min at 550g ; 16 °C) and washed in 1 mL cold (4 °C) MAX Efficiency buffer (invitrogen). The pellet was then re-suspended in 100 µL cold (4 °C) MAX Efficiency buffer and 15 µg linearized plasmid DNA was added before loading into an 0.2mm gap electroporation cuvette (Sigma Z706086).

Electroporation was carried out using the following settings (Yamano et al. 2013) in a Nepagene NEPA21 instrument. Immediately following electroporation, algae were removed from the cuvette and incubated in 40 mL AF6 media at 16 °C. Poring : 250V ; 2 pulses ; *Pulse length 8 msec* ; *Decay rate 40 %* ; *Pulse interval 50 msec* Tranfert : 20V ; 10 Pulses ; *Pulse length 50 msec* ; *Decay rate 40 %* ; *Pulse interval 50 msec*

#### RNA extraction

RNA extractions were performed using the RNeasy Plant Mini Kit (Qiagen). 5 mL of liquid culture at maximum density were pelleted at 550g for 5 minutes and cell pellets were re-suspended in RNA lysis buffer solution supplemented with B-mercaptoethanol. The manufacturer’s protocol was then followed to extract RNA from *Oophilla* cells.

#### RT-qPCR

To produce cDNA from total RNA, QuantiNova Reverse Transcription Kit (Qiagen) was used with 1 µg of total RNA for each sample. Following the RT reaction, cDNAs were diluted tenfold in water (QuantiNova SYBR Green PCR, Qiagen). 1 µL of 10fold diluted cDNA was loaded in duplicate on a MIC PCR (BioMolecularSystem) instrument and the following cycle program was executed: 95 °C for 3 min ; 35x[95 °C for 5 sec, 60 °C for 15 sec] ; melting curve (75 to 90°C ; 0.3°C/sec). qPCR mix consisted in : Master mix 1X final, primers 0.7 µM final and 1 µL 10-fold diluted cDNA. Cq data was exported and results were expressed using the 2^-ΔCt formula.

#### Primer list

##### Reference genes

p232_Ooph_RACK1_L_3 CGCACAGCCAGTAGCGGT

p233_Ooph_RACK1_R_3 GGACCTGGCTGAGGGCAA

p234_Ooph_YPTC1_L_4 TTGCGGATGACACCTACACG

p235_Ooph_YPTC1_R_4 TGGTCCTGAATCGTTCCTGC

#### Transgene specific primers

p189_aphviiiUTR_qF tatcggaggaaaagctggcg

p190_aphviiiUTR_qR gatcccaacgtccacactgt

#### Western blot

Total proteins were extracted in a buffer containing 25 mM Tris pH 7.5, 12 mM MgCl2, 15 mM EDTA, 75 mM NaCl and 0.1% tween 20. In addition, Protease and Phosphatase Inhibitor Mini Tablets (Pierce) and 1 mM DTT were added before use. Protein concentration was determined by a Bradford assay (Quick start Bradford Protein Assay, Biorad) and 0.75 µg/µL samples were boiled in SDS-loading buffer. 6X SDS-loading buffer composition is 30 % Glycerol, 9 % SDS, 0.6M DTT, 0.012% Bromophenol blue 0.35M Tris HCl pH 6.8 (adapted from Laemmli 1970).

Proteins were resolved by 12% SDS-PAGE at constant amperage of 20 mA per gel in tris-glycine running buffer (25 mM Tris, 0.192 M glycine, 0.1% SDS, pH 8.3). Proteins from the gel were then transferred to a nitrocellulose membrane ((0.45 µm) from Amersham Protran) in a wet tank system (Mini Trans-Blot, Biorad) at 100V for one hour. Following transfer, membranes were blocked for 45 minutes in PBS-0.1%Tween20 (PBS-T)-5% dry milk. For protein detection, membranes were incubated overnight in PBS-T-5% dry milk with anti-FLAG antibodies (Sigma F1804) at 1:10000 dilution. Acrylamide gels were stained after blotting in coomassie blue for protein loading controls.

Blots were washed three times for 15 minutes with PBS-T and probed 45 minutes with secondary antibodies (rabbit anti-mouse (Sigma A4312), 1:10000 dilution) conjugated to alkaline phosphatase in 1x PBS-T 5% dry milk. After three washes in PBS-T, the blot was washed in alkaline phosphatase buffer (100 mM Tris-HCl pH 9.0, 150 mM NaCl, 1 mM MgCl2) for 10 minutes and BCIP/NBT solution (Promega PAS3771) was added following manufacturer instructions.

#### Flow cytometry

Cells were filtered through a 40 µm mesh and loaded in a ZE5 Cell Analyzer flow cytometer (Bio-Rad). Data was collected for forward scatter and side scatter using the 488 nm laser. GFP fluorescence was measured using excitation from a 488 nm laser and a 525/35 nm band-pass emission filter. Algal chlorophyll fluorescence was measured using a 488 nm laser for excitation with a 692/80 nm band-pass emission filter. Flow cytometry data was analyzed in R v4.1.0 (R Core Team 2013) using packages flowCore v2.4.0 (Ellis et al. 2022), ggcyto v1.20.0 (Van P et al. 2018), flowClust v3.30.0 (Lo et al. 2009), and flowStats v4.4.0 (Hahne et al. 2021). Algal cells were identified using automated clustering (tmixFilter) in the side scatter and chlorophyll fluorescence channels, and the cell population was analyzed for GFP fluorescence levels.

#### Optical microscopy

Live cells were observed using a Leica DMi8 inverted epi-fluorescence microscope. GFP was visualized under a 480 nm excitation laser and a 510 nm emission filter, and captured with a DFC9000 sCMOS camera.

## Acknowledgements

We thank Dr. Cory Bishop for sharing his unpublished draft genomes of the algae *O. amblystomatis, Chlamydomonas nasuta, Chlamydomonas moewusii*, and *Chlamydomonas pseudogloegama*. Comparative analyses of those genomes enabled promoter region selection.

